# Assessing the optimal frequency of early parasitoid releases in an apple orchard to control *Dysaphis plantaginea:* a proof of concept study

**DOI:** 10.1101/2021.09.20.461076

**Authors:** L. Ferrais, K. Tougeron, P. Gardin, T. Hance

**Affiliations:** Earth and Life Institute, Ecology and Biodiversity, Université catholique de Louvain, croix-du-Sud 4-5, 1348 Louvain-la-Neuve, Belgium

**Keywords:** biological pest control, natural enemies, temperature, mass release, release timing

## Abstract

Alternative measures to pesticides to control the rosy apple aphid *Dysaphis plantaginea* are being developed. Naturally occurring predators and parasitoids often fail to reduce aphid abundance below the economic threshold in orchards, because they are active too late after the aphid first infestation. We tested the efficiency of mass release of two parasitoid species, *Aphidius matricariae* and *Ephedrus cerasicola*, early in the season to match the presence of aphid fundatrix (sensitive stages). In this trial focusing on an organic apple orchard, three releases were done either every week or every two weeks to test the effect of the release frequency, during two consecutive years. The number of aphid colonies and aphid number per tree were monitored from late March to late May. Degree-days necessary for parasitoid emergence in the field after release were calculated. We show that a sufficient level of aphid control by parasitoids is reached during the first month of the survey, but control mostly fails during the second part of the monitoring session, for both release treatments, and compared to the neem oil control treatment. The relative effects of release frequencies were different between years probably because of interannual differences in aphid population dynamics and initial infestation in orchards. The field survey and the degree-day model suggest that parasitoid releases, at either frequency, are promising candidates for biological control of the rosy aphid, although the method still needs proper calibration. This conclusion needs to be reinforced by repeating the study in more orchards, but our case study lays the first empirical basis that will help to develop future control methods of aphids by parasitoid releases in apple orchards. We argue that releases should be done one to two weeks before first aphid detection to account for long development times of parasitoids at relatively low temperatures.

## Introduction

Apple production has a huge economic value worldwide; it represents one of the most important fruit crop and contributes to almost 35% of European orchards (FAO 2018). However, the sustainability of apple production is becoming controversial since it relies on a heavy use of pesticides to control different kinds of pests (Reganold et al. 2001; Simon et al. 2011). As a result of frequent insecticide applications, resistant pest strains have been reported for most registered chemicals (Varela et al. 1993). Development of Integrated Pest Management (IPM) approaches have been undertaken since the 1970’s in an attempt to limit the use of pesticides (Blommers 1994; Cross et al. 2015; Heijne et al. 2015). Within the current framework of new policies regarding chemicals (e.g., neonicotinoid ban in some countries), concerns on health- and environment-damaging risks, and public growing demand for organic food production, there is an urgent need to design innovative solutions for pesticide-free orchard protection (Zehnder et al. 2007; Simon et al. 2017). Fortunately, only a few arthropod species present on apple trees are actual major pests, but they are causing damage and yield loss even at low population densities (Cross et al. 2015).

In apple orchards of Europe and North America, one of these major insect pests is the rosy apple aphid *Dysaphis plantaginea* (Hemiptera: Aphidinae), causing leaf-rolling and fruit deformation (Wilkaniec 1993; Brown and Mathews 2007), and significant yield losses when uncontrolled (Dib et al. 2010). The aphid is holocyclic, with two successive host plants: apple and plantain (*Plantago lanceolata*) (Blommers et al. 2004). In early spring, fundatrix females hatch from diapausing eggs deposited around the buds. The presence of those fundatrix coincides with bug break, which is followed by several generations of wingless parthenogenetic viviparous females in the spring and early summer, responsible for most of the damages on apple trees. By the time trees bloom, the leaves begin to curl, providing protection to aphids (Brown and Mathews 2007). Winged morphs are eventually produced and migrate to plantain in summer. Aphids return to apple in early fall and produce the sexual generation that will lay overwintering eggs (Blommers et al. 2004). One stage of the life cycle is particularly relevant as a target for control methods: fundatrix aphids, because they are the starting point of an exponential and massive parthenogenetic reproduction in early spring, which will be damaging trees. Additionally, they hatch from eggs at relatively low abundances (depending on sexual aphid abundance in fall).

The search for environmental-friendly and economically sustainable solutions for pest control in IPM has stimulated interest towards *D. plantaginea* natural enemies. Aphid midges, hoverflies, earwigs, predatory mites, ladybugs and spiders are among the most abundant predatory arthropods in Europe and North America reducing populations of the rosy aphid (Miñarro et al. 2005; Brown and Mathews 2007; Cross et al. 2015). Parasitic wasps, mostly Braconidae and Mymaridae, are also slowing aphid population growth (Bribosia et al. 2005; Peusens et al. 2006; Dib et al. 2012), although their efficiency is higher when paired with the generalist predator guild (Gontijo et al. 2015). However, most of these natural enemies occur too late in the aphid’s lifecycle to exert strong regulating effects on these already large colonies formed by fundatrix aphids (Miñarro et al. 2005; Bribosia et al. 2005). For this reason, in organic production, naturally occurring predators and parasitoids often fail to reduce aphid abundance below the economic threshold (Brown and Mathews 2007; Dib et al. 2010), especially in years of high aphid density (Wyss et al. 1999). Therefore, augmentative releases of beneficial insects in early spring have been proposed to complement the impact of the naturally occurring natural enemies of aphids (Kehrli and Wyss 2001; Dib et al. 2016).

Parasitoid cocktails are used in inundative biological control strategies in several crop systems, because each species usually complements the others on their host spectrum, and on behavioral and physiological specificities (Dassonville et al. 2012; Boivin et al. 2012). A cocktail of two solitary parasitoid species, *Aphidius matricariae* and *Ephedrus cerasicola* (Hymenoptera: Braconidae), was developed in previous projects to control *D. plantaginea* in apple orchards (Dumont et al. 2011; Boivin et al. 2012; Nicolas et al. 2013). These two parasitoid species are known to parasitize and properly develop in the rosy apple aphid (Hullé et al. 2006). Although this cocktail was promising in laboratory studies, preliminary experimental studies still showed poor control of the rosy aphid, probably because of mistiming with aphid first infestation, and of high heterogeneity of parasitoid establishment after introduction (Hance et al. 2017). The aim of the present study was to perform a first exploratory trial of parasitoid releases in an orchard— which has never been done before—from which future studies can build on to develop biological control methods against the rosy apple aphid by early mass releases of aphid parasitoids to match aphids’ sensitive life-stage. Our goal was first to determine the optimum frequency of parasitoid releases that would allow sufficient regulation of fundatrix aphids in early spring, and secondly to obtain degree-days necessary for parasitoid emergence in the field after release. Release frequency is a major issue, because a trade-off has to be found between continuous presence of parasitoids in orchards over the time window of aphid presence, their overall efficiency, and the importance of reducing intraguild competition and superparasitism. For example, *A. matricariae* and *E. cerasicola* are known to compete through multiparasitism on the aphid *Myzus persicae* (Hågvar 1988). Based on laboratory results, we hypothesized that every-two-weeks releases would be optimal, because it would limit parasitoid generation overlapping and allow spreading parasitoid presence over a long period in orchards.

## Material and Methods

### Study area and release protocol

In order to determine the optimal parasitoid release time and frequency, corresponding to the time when more than half of the aphid fundatrix emerged, we used a model based on the calculation of accumulated degree-days (DDs) by aphid eggs in winter. According to this model, *D. plantaginea* eggs need an accumulation of 110 to 230 DD, with an average of 153 DD to hatch (Demeter et al., unpublished data). Similar studies in the literature set the average at 140 DD (110 to 180 DD) (Graf et al. 2006). The value of degree days accumulated in a day by aphid eggs is calculated by summing the DDs of the 24 hours of the day, using the following formula:

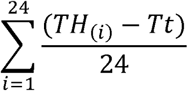

Aphid eggs are accumulating DDs as soon as the outside temperature exceeds 4°C, which represents their developmental temperature threshold T*t* (Graf et al. 2006). For each hour of the day, the threshold temperature T*t* is subtracted from the temperature of the studied hour TH_(i)_. The result is then divided by 24 to obtain a mean hourly value. Degree-day calculation begins on the 15th January, the date at which eggs are known to emerge from 90 days of diapause (Graf et al. 2006). Then, DDs are summed up for each day from the 15th January until the day at which a total of 153 DD is obtained.

Our experiment was carried out in spring from April 11th to May 23rd, 2018, and from March 28th to June 21st, 2019, in an organic commercial apple orchard located in Fleurus, Belgium (50.47°N, 4.54°E). The protocol was conducted in two rows formed of 108 apple trees (*v*. Natyra) at the outermost edge of the orchard. Rows of apple trees were separated into three groups corresponding to the three treatments applied; parasitoids released weekly, parasitoids released every two weeks, and a control treatment consisting in neem oil (a standard organic treatment that was applied to the rest of the orchard one time before blooming and another time after if aphids persist) (**Figure 1**). Each zone was similar in all respects to the others. It was unfortunately not possible to maintain an untreated area within the commercial apple orchard, as growers generally do not want to take the risk of uncontrolled aphid outbreaks.

**Figure 1:**
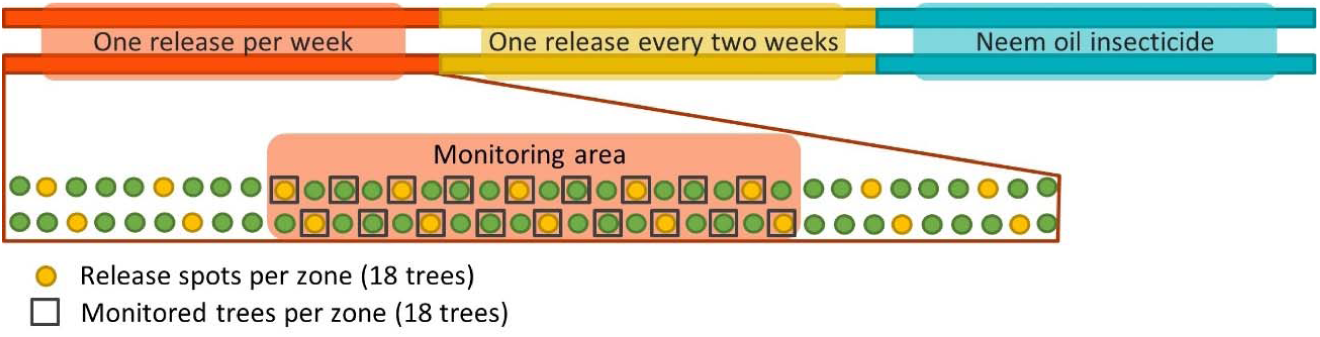
Release and monitored trees for 2018. For 2019, zones where the “one release per week” and “one release every two weeks” treatments were applied in 2018 were switched. Each zone was similar in all respects to the others. For each tree line, parasitoids were released one tree out of four and monitoring was done every two trees in the central area.

The mix of parasitoids species was provided by Viridaxis SA (Belgium) in cardboard tubes, each containing 450 *A. matricariae* and 530 *E. cerasicola* close to their final development stage. We managed to obtain a density of parasitoid per release tree of 114 *A. matricariae* and 135 *E. cerasicola* (Demeter et al., unpublished data). Parasitoids were released on one out of four trees (total of 36 trees) at three occurrences. The releases of 2018 were conducted on the 11th, the 18th and the 25th of April (one release per week treatment), and the 11th, the 25th of April and the 09th of May (one release every two weeks treatment). For 2019, parasitoids were released on the 28th of March, the 04th and the 10th of April (one release per week treatment), and on the 28th of March, the 10th and the 24th of April (one release every two weeks treatment). Zones applied with parasitoid releases every week or every two weeks were switched between 2018 and 2019 to control for any effect of the position on the tree row.

### Monitoring protocol

Within each release zone, we delimited a central area of 36 trees (18 trees per row, in order to avoid a border effect (Holland and Fahrig 2000)), where aphid populations were monitored weekly (**Figure 1**). The number of aphid colonies (i.e., buds infested with rosy apple aphids) were counted, as well as the number of aphids per colony for 10 randomly selected colonies per monitoring zone (5 on each row). Colonies were marked to allow monitoring of the same colonies over weeks. In the first few weeks of monitoring, fundatrixes in the marked colonies may move around the tree or disappear. Therefore, when an aphid colony counted two or less aphids, another colony on the same tree was marked to replace it, in order to have continuous observations across the season.

### Parasitoid emergence and degree-day calculation

To study the emergence of parasitoids, cardboard tubes containing mummies were attached to cylindrical plastic tubes on apple trees and closed with a piece of tulle. Three tubes containing 50 *A. matricariae* and three tubes containing 50 *E. cerasicola* mummies were followed at each parasitoid release date, except for the first week because of technical problems with the sleeves-cage. A fourth replicate was thus done to have a total of three replicates. Parasitoid emergence was counted three days per week, on Mondays, Wednesdays, and Fridays. We measured the emergence rate (number of emerged parasitoids / total number of mummies) and the sex-ratio (number of males / total number of emerged parasitoids). Using the formula used for rosy apple aphid eggs, the number of DD needed for development from mummy to adult was calculated for the two species of parasitoids. Temperatures above 2.00°C for *A. matricariae* and above 6.83°C for *E. cerasicola* were taken as baselines (Cohen and Mackauer 1987; Colinet and Hance 2010). The calculation of DD was done from the first day of release until the day we obtained 50% of parasitoid emergence.

### Statistical analyses

Data collected from the two years were analyzed separately because there were variations in the initial infestation date of *D. plantaginea* and in climatic conditions. The number of aphid colonies per tree and the mean number of aphids per tree (mean aphid number per colony / number of colonies per tree) were fitted to negative binomial Generalized Linear Mixed Models (GLMMs), using the *glmmTMB* package in R (Magnusson et al. 2017). In 2019, most of the market colonies had disappeared at the three last dates of survey so aphid number per tree could not be calculated properly. Therefore, these three dates were excluded for the dataset of the aphid number per tree. We used the date, the release treatment (three levels: every week, every two weeks, neem oil), and the interaction between these factors as explanatory variables in our models. The identity of the apple tree (N = 54) was used as a random effect in the GLMMs to correct for pseudoreplication and repeated measures. A preliminary analysis was conducted by adding a “release tree” factor in the models (two levels: tree is a parasitoid release point, or not). We showed that there were no differences in the number of aphids (2018: χ^2 =^ 0.38, p = 0.5, 2019: χ^2 =^ 0.68, p = 0.4), and in the number of colonies per tree (2018: χ^2 =^ 0.01, p = 0.9, 2019: χ ^2 =^ 0.27, p = 0.6) between trees on which parasitoids were released and other trees, so the “release tree” factor was removed from the models for further analyses. Model selection was done with the function *Anova* from the package *car* (Fox and Weisberg 2011); fixed effects were kept in the model when p<0.05. Conditional R-squared for GLMMs (proportion of total variance explained through both fixed and random effects) were calculated using the package *MuMIn*. Multiple comparisons among levels of significant variables were performed using the *glht* function from the *multcomp* package (Hothorn et al. 2008). Model goodness of fit and dispersion parameters were verified using the *DHARMa* package (Hartig 2020), and no significant problems were detected. Finally, an unpaired two-samples t-test was used to compare the mean of the sex ratio between the two parasitoid species (normal distributions with equal variances). Emergence rates were compared between parasitoid species using a Welch t-test (to account for normal distributions but unequal variances). All analyses were carried out using R (R Core Team 2020).

## Results

### Field experiment of 2018

The date, the treatment, and their interaction showed significant effects on the mean number of aphids per tree, and on the number of colonies per tree (with a marginally non-significant effect of the interaction term for this parameter) (**Table 1**). In 2018, the initial infestation of *D. plantaginea* was weak, with close-to-zero aphid colonies and aphid per tree on the 11th and 18th of April (**Figure 2a, 2b**). After that, the colony number and aphid number per tree increased, especially for the zone where releases were done every week. Overall, the zone where the three parasitoid releases have been done every week showed a higher mean (± SE) of aphid colonies than both other zones (2.40 ± 0.45 *vs*. 0.53 ± 0.23 for the two-weeks-release treatment (glht, z=3.2, p<0.01), and 0.54 ± 0.12 for the control zone (z=2.9, p<0.01)), as well as a higher mean number of aphids per tree (265 ± 78 *vs*. 44 ± 29 for the two-weeks-release treatment (z=3.8, p<0.001), and 12 ± 3 for the control zone (z=4.9, p<0.001)). These differences among treatments were mainly visible at the last three monitoring dates.

**Table 1:**
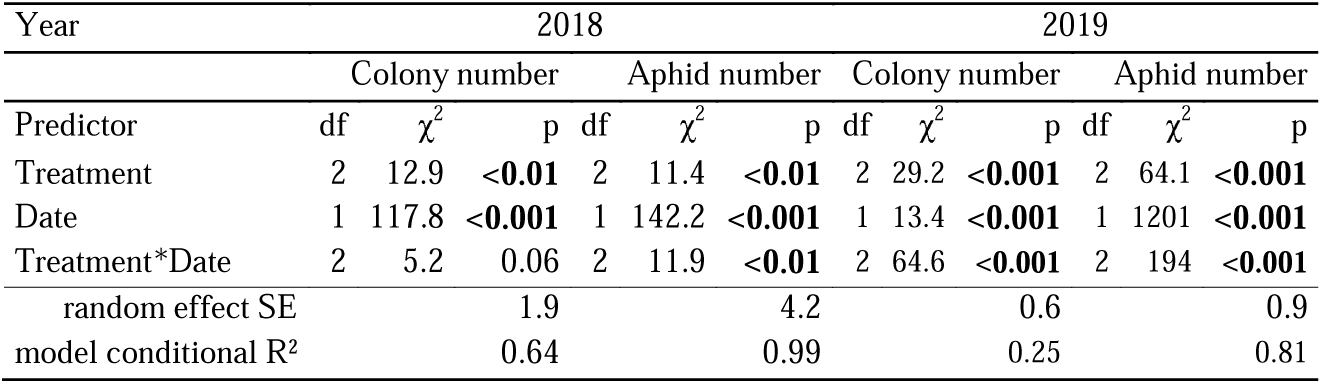
Statistical results (type II Anova after a GLMM) for *Dysaphis plantaginea* colony numbers per tree and the aphid numbers per tree in 2018 and 2019 surveys. Significant results are in bold.

| Year | 2018 |  |  |  |  |  | 2019 |  |  |  |  |  |
| --- | --- | --- | --- | --- | --- | --- | --- | --- | --- | --- | --- | --- |
|  | Colony number |  |  | Aphid number |  |  | Colony number |  |  | Aphid number |  |  |
| Predictor | df | $\chi^2$ | p | df | $\chi^2$ | p | df | $\chi^2$ | p | df | $\chi^2$ | p |
| Treatment | 2 | 12.9 | **<0.01** | 2 | 11.4 | **<0.01** | 2 | 29.2 | **<0.001** | 2 | 64.1 | **<0.001** |
| Date | 1 | 117.8 | **<0.001** | 1 | 142.2 | **<0.001** | 1 | 13.4 | **<0.001** | 1 | 1201 | **<0.001** |
| Treatment*Date | 2 | 5.2 | 0.06 | 2 | 11.9 | **<0.01** | 2 | 64.6 | **<0.001** | 2 | 194 | **<0.001** |
| random effect SE |  |  | 1.9 |  |  | 4.2 |  |  | 0.6 |  |  | 0.9 |
| model conditional R<sup>2</sup> |  |  | 0.64 |  |  | 0.99 |  |  | 0.25 |  |  | 0.81 |

**Figure 2:**
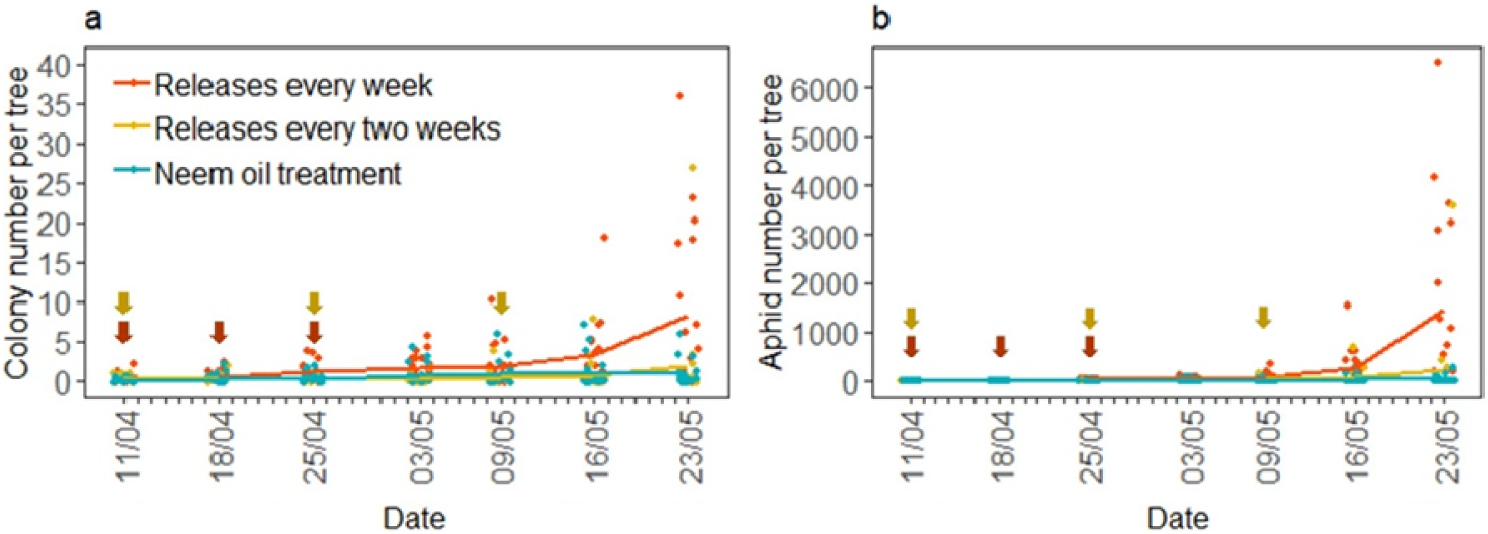
(a) Number of colonies and (b) number of aphids per tree according to the three treatments, for all dates in 2018. Lines correspond to mean values. Red arrows represent releases carried out every week, and yellow arrows represent releases carried out every two weeks.

### Field experiment of 2019

In 2019, the early infestation of *D. plantaginea* was greater than in 2018, with a mean of 13 ± 1 colonies and 61 ± 10 aphids per tree, respectively. Both the number of aphid colonies per tree and the number of aphids per tree were explained by the treatment, the date and their interaction (**Table 1**). After the first monitoring date, there was a slight decrease in the number of colonies and of aphids per tree, but these numbers increased again to reach a peak on the 29th May (**Figure 3a, 3b**). Over the entire monitoring season, the aphid infestation was higher in the zone where the three parasitoid releases were done every two weeks (**Figure 3a, 3b**) for the number of colonies (18.23 ± 1.47 *vs*. 8.94 ± 0.84 for the one-week treatment (z=3.8, p<0.001), and 5.60 ± 0.29 for the control treatment (z=4.9, p<0.001)). It was also the case for the number of aphids per tree (2137 ± 353 *vs*. 1010 ± 192 for the one-week treatment (z=2.8, p<0.05), and 50 ± 4 for the control treatment (z=9.3, p<0.001), which itself differed from the one-week treatment (z=6.0, p<0.001)). However, these differences among treatments were not observed at every date; the mean numbers of colonies per tree were similar among treatments on the 7th of May and at the end of the monitoring period, but were different at the very beginning of the monitoring and around the infection peak. Similarly, the mean number of aphids differed among treatments at the beginning and the end of the monitoring session but was similar around May 7th. However, we observed that the trend of aphid density evolution across the monitoring season remained the same between both parasitoid release treatments.

**Figure 3:**
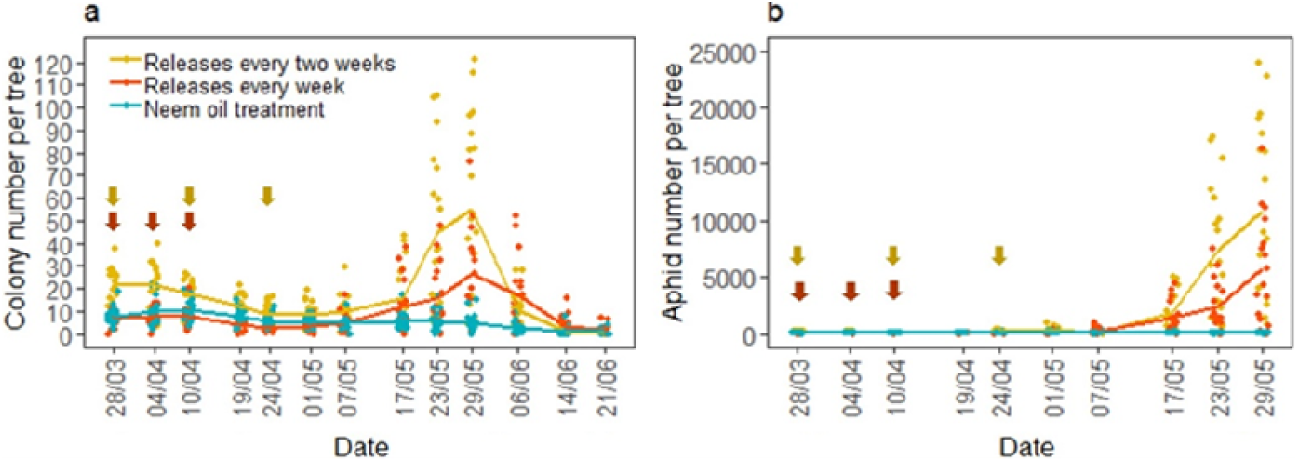
(a) Number of colonies and (b) number of aphids per tree according to the three treatments, for all dates in 2019. Lines correspond to mean values. Red arrows represent releases carried out every week, yellow ones, releases carry out every two weeks and blue arrows represent the date of neem oil treatment.

### Parasitoid emergence and degree-day calculation

Parasitoid emergence rate ranged between 48.3% and 90.4% for *E. cerasicola* and 76.9% and 100% for *A. matricariae*, with a different mean (±SE) of 66.3 ± 4.2% and 88.2 ± 1.9%, respectively (**Table S1**, *t* = 4.7, *p* <0.001). The sex ratio ranged between 0.47 and 0.70 for *E. cerasicola* and 0.35 and 0.77 for *A. matricariae*, with a similar mean (±SE) of 0.60 ± 0.02 and 0.52 ± 0.04, respectively (**Table S1**, *t* = -1.5, *p*= 0.15).

For each species and within each replicate, the date at which 50% of parasitoids did emerge were similar for the three cages of a given replicate, but was different between all the replicates and between species (**Table S1**). *Aphidius matricariae* took an average of 8 days to emerge (min-max 3-12 days) and *E. cerasicola* took an average of 16 days (min-max 9-21 days). Degree-days required for emergence were similar among cages of each replicate, for each species, but varied among replicates and species. Mean degree-day value for *A. matricariae* was 69.47 (min-max 51.43-98.65) and for *E. cerasicola* 146.40 (min-max 112.60-177.38). Degree-days were the lowest for the second replicate for both species, and the highest for the last replicate for *A. matricariae* and the first replicate for *E. cerasicola*.

Based on the degree-day values, we calculated the date at which 50% of parasitoids from each species should have emerged in the field for the 2019 year. It allowed mapping the potential presence time of parasitoids in the field with the presence of aphids, depending on temperatures, parasitoid release dates and release frequency (**Figure 4**). Parasitoid emergence was less staggered over time when temperatures were higher, and *E. cerasicola* took overall more time to emerge than *A. matricariae*, even if they were released at the same time. When releases were done every week, the three first emergence waves of *E. cerasicola* occurred at the same time as the increase of both aphid number and colony number per tree. Concerning *A. matricariae*, the two first emergence waves happened early, and the third wave occurred just before the rise of aphid populations. When releases were done every two weeks, *A. matricariae* from the first wave had emerged simultaneously with the decrease of aphid number per tree and colony number (from April 4th to April 10th). The third parasitoid wave emerged just before aphid number rose. The fourth wave emerged during a decrease of aphid number, between May 1st and May 7th. For *E. cerasicola*, the first and third emergence waves occurred during a slight increase of aphid number, which then declined quickly. The fourth wave happened just before a strong rise of aphids (from the 08th to the 17th of May).

**Figure 4:**
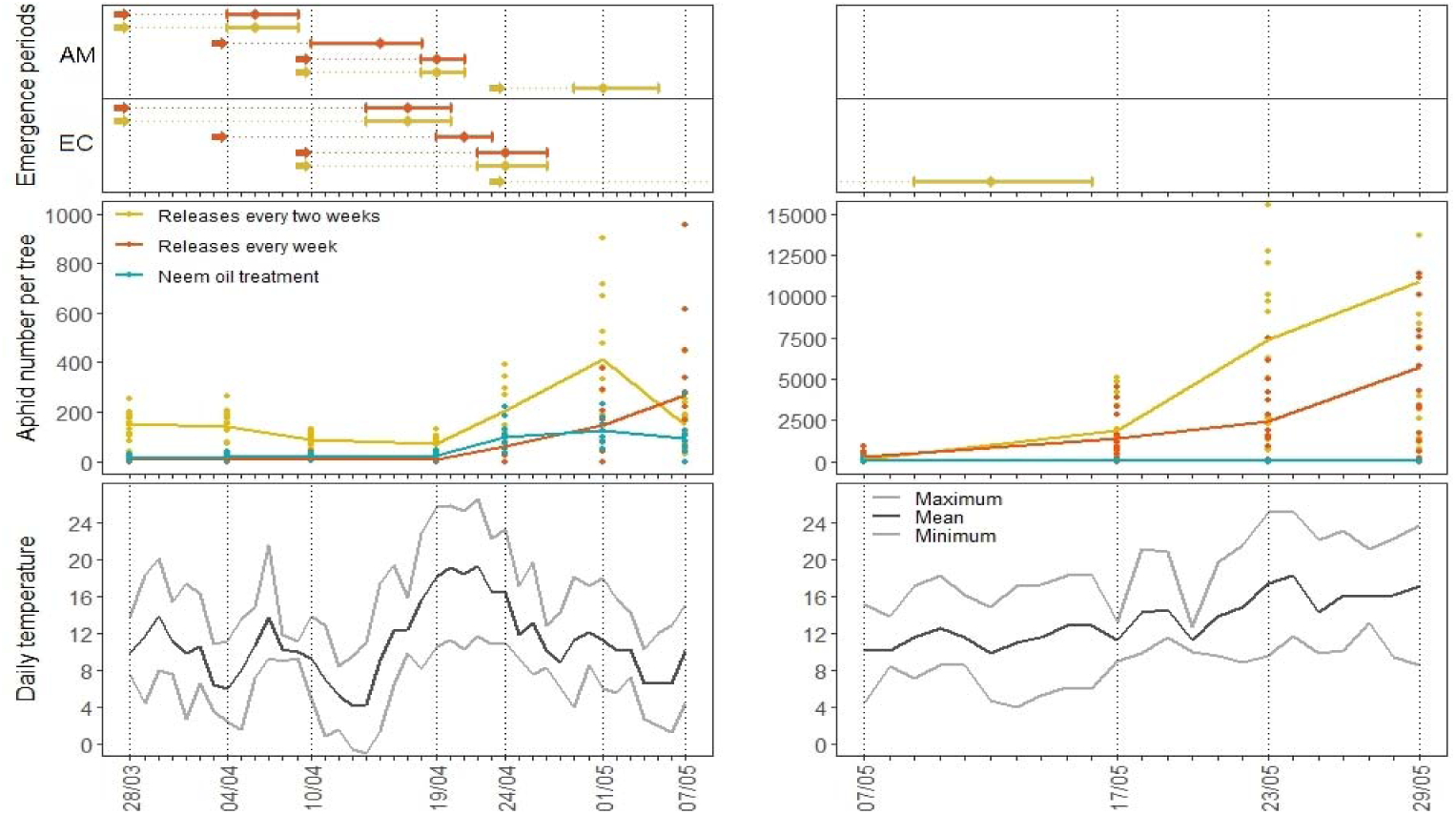
Mean number of aphids per tree according to the three parasitoid release treatments from March 28^th^ to May 7^th^ (left panel) and from May 7^th^ to May 29^th^ (right panel), 2019. Bottom graphs show the minimum, mean, and maximum daily temperatures (°C). Dotted lines correspond to the dates of aphid survey in the field. Top graphs show the predicted presence (emergence period) of parasitoids in the field for both species, according to calculated degree-days. (For each species (AM: *Aphidius matricariae*, EC: *Ephedrus cerasicola*), arrows represent release dates: every week (red) or every two weeks (yellow). The lines represent min-max range of parasitoid predicted presence, and the points represent the mean emergence time.

## Discussion

Infestation of the orchard by *D. plantaginea* (as defined by the number of aphid colonies and mean aphid numbers per tree) was, as expected, highly dependent on the date, but also varied depending on the parasitoid release treatment, for both monitoring years. Of course, our case study was conducted as an exploratory assay in one apple orchard over two years to test the validity of the method in terms of biological control, so we discuss our results being aware that it is not yet possible to fully generalize the conclusions. Nevertheless, and keeping in mind this caveat, our experience provides first in-field empirical results to continue developing programs based on parasitoid releases to control the rosy apple aphid in orchards, and we hope it can lay the foundations for future studies. In addition, results on the degree-days necessary for parasitoid emergence after release can help constructing phenological models and adjusting the release timing of both species.

The infestation pattern and dynamics of rosy apple aphids were quite different from one year to another. For example, the initial quantity of aphids was lower in 2018 than in 2019, which may have governed the infestation dynamics of the rest of the monitoring season. In 2018, regardless of the treatment, the number of colonies and aphids per tree slowly increased until the end of the monitoring session to reach a peak. In 2019, aphid infestation slightly decreased during the first month of the survey before reaching a peak at around the same date as in 2018 (end of May). As colony monitoring has continued in 2019 until the end of June, we observed after the peak a sharp decrease in aphid densities, probably due to migration of rosy apple aphids toward their secondary host plant (Blommers et al. 2004). Overall, our results concur to the idea that efficient biological control strategies against *D. plantaginea* should be set up as early as March, when aphid densities are still low enough.

For both study years, the zone on which neem oil was applied (control treatment) showed relatively good control of rosy apple aphids. Neem has been tested against a large number of insect pests, and have shown some efficacy against apple aphid species (Schulz et al. 1997; Kraiss and Cullen 2008; Peck 2010). However, neem products are relatively expensive and several applications at short intervals are required for pest control (Peck 2010), so alternative solutions such as parasitoid releases are needed. In addition, neem oil could have non-target effects because it is moderately toxic for beneficial insects such as pollinators, although less harmful than non-natural products (Peck 2010; Aliakbarpour et al. 2011). In our study, the neem oil treatment served as a control and was in some instances not sufficient to fully regulate aphid populations, especially early in the spring when apple trees are sensitive to aphid attacks.

Concerning parasitoid action on aphid populations, we first showed that releases occurring every two weeks were as effective as the treatment with the neem oil, in 2018. Aphid populations were effectively controlled until the end of the monitoring season, whereas they were consistently larger, and they increased faster in the every-week-release treatment. In 2019, initial aphid infestation was higher in the zone with parasitoid release done every two weeks, but this treatment still showed a similar trend as other treatments until May 17th, suggesting relatively good aphid control. One could speculate that releases done every two weeks are more efficient when initial aphid densities are low, and releases done every week are optimal when early-season aphid densities are high. However, if parasitoid releases seemed as effective as the neem oil treatment in early season, it was less effective later on, especially with the two-weeks-release treatment. It is therefore clear that parasitoid releases did not allow proper control of the infestation of *D. plantaginea*, because both release frequency treatments showed less efficiency than the neem oil treatment over the entire monitoring session in the orchard we monitored.

Our results indicate that parasitoid releases were not fully effective to control rosy apple aphids in this orchard, especially when the initial infestation was strong, such as in 2019. It is likely that the number of colonies and aphids depend on spatio-temporal constraints in the orchard, that affect control efficiency by parasitoids. Indeed, spatial distribution of *D. plantaginea* in orchards is probably not random and may depend on multiple factors such as the presence of alternate host plants, tree age, planting densities and the surrounding habitat (Kozar et al. 1994; Brown and Myers 2010). To the same extent, parasitoid presence and persistence in orchards are influenced by local- and landscape-scale environmental conditions and practices (Jahnke et al. 2008; Maalouly et al. 2013; Kishinevsky et al. 2017). In addition, we encourage future studies to consider as an additional variable the phenology of hyperparasitoids that tend to be increasingly present early in the season, especially in warm years, and which could impair biological control in numerous crop systems (Tougeron and Tena 2019).

The density-dependence control of aphids still needs to be assessed in orchard conditions, because there might not be a clear functional response between the number of parasitoids that are released and aphid densities (Nicolas et al. 2013; Hance et al. 2017). Indeed, parasitoids may disperse out of the release areas and out of the orchard, although the vast majority of parasitism occurs close to the release points (Dumont et al. 2011; Boivin et al. 2012; Zappala et al. 2012; Nicolas et al. 2013). Therefore, there is a need to find a balance between dispersion within the orchard, to limit the number of release points and maximize homogeneous establishment, and dispersion out of the orchard that would be pure waste in terms of biological control performance. Same issue is arising in other cultures for mass release of parasitoids (Van Lenteren 2000). Therefore, an intermediate level of dispersal may maximize establishment and guarantee an effective distribution of parasitoids (Heimpel and Asplen 2011; Zappala et al. 2012). Infield management such as the provision of alternative food-sources within tree rows (e.g., flower strips) or protection against wind and deleterious climatic conditions may both improve the dispersal ability of the natural enemy released within the orchard, and prevent them from dispersing out of the orchard.

Both parasitoid species used for the field experiments differed on their emergence rates with a better success of *A. matricariae* (88.2%) than of *E. cerasicola* (66.3%). Many factors could explain these contrasts, such as for example differences in thermal tolerance capacities between both species (e.g., Le Lann et al. 2011), leading to increased mortality in *E. cerasicol*a during early cold snaps of the monitoring season. Increased mortality in *E. cerasicola* could also be due longer exposure to hazardous conditions, because it takes 5 days at 17°C for this species to emerge from the mummy, while it takes 3 days for *A. matricariae* (Viridaxis SA, personal communication). The sex ratio was similar between both species and remained quite high with more males produced for both species, which in unusual in natural populations of parasitoid wasps, but expected in commercial strains (Heimpel and Lundgren 2000). It means that of the total number of parasitoids released in the orchard, more than a half were males, not actively researching rosy apple aphid hosts. Unbalanced sex ratio can be due to differential mortality of parasitoids of each sex, as it can happen that fewer pre-imago females survive detrimental conditions (cold, drought, bad host quality, pathogen infection, exposure to pesticide, etc.), compared to males (Langer et al. 2004; Joseph et al. 2011; Tougeron et al. 2020).

In inundative biological control strategies, one seeks to maximize parasitoid presence over time in the target crops, and to match it with the presence of target pests, which can be achieved by multiplying release events, but also by taking advantage of differences in development time among released species (Van Lenteren 2000). Degree-days obtained for both parasitoid species are good indicators of the ideal time to put parasitoid cardboards in the orchard. At low temperatures in outdoor conditions at the beginning of the monitoring season, parasitoid emergence took a long time (9 to 20 days after release), leading to parasitoid presence late after aphid appearance. In addition, degree-days required for *E. cerasicola* to emerge were almost two times higher than for *A. matricariae*. To improve acute parasitism pressure on *D. plantaginea* over a short period of time (e.g., when fundatrix aphids are present), we would recommend a desynchronized release of the two species, with *E. cerasicola* releases done a week before releases of *A. matricariae*. On one hand, differences in emergence time between both parasitoid species might be an asset for biological control, because it allows staggering their presence in orchards. On the other hand, each species might be as efficient alone as it is in combination with the other, given their emergence timing and emergence rate, and because no study has yet assessed the potential of intraguild competition between both species in orchard conditions. The rationale of using both species in the release mix is thus still to be investigated.

To conclude, parasitoid releases made at both one-week and two-week frequencies showed some effects early in the season on decreasing, or at least stabilizing aphid and colony numbers in the apple orchard. Unfortunately, parasitoid releases did not seem as efficient as the control neem oil treatment to maintain aphid populations under an acceptable threshold over the entire apple tree sensitive period. We suggest that parasitoid releases were not done early enough to ensure parasitoid-host synchrony, and cardboards should be placed in orchards around 10 days (one to two weeks) before the first appearance of fundatrix aphid on apple trees to account for the long development time of parasitoids in the early season. However, the main caveat to this solution are low temperatures that can have a negative impact on parasitoid emergence rates, fitness, and overall efficiency. Releases could also be done over a longer period of time and/or with increased release frequencies. On this point however, we stress that such inundative release methods are time-consuming to apply in the fields, and still presently represent an important cost for apple producers. Cheaper solutions for parasitoid mass rearing are still to be developed (Van Lenteren 2000; Levie et al. 2005), with strains active at low temperatures and a more balanced sex-ratio (Van Lenteren 2000; Boivin et al. 2012).

## Disclosure statement

We have no conflict of interest to declare.

## Acknowledgements

This article is part of the ERAnet C-IPM project API-Tree. LF and PG were supported by the API-Tree project. KT was supported by the F.R.S.-FNRS. We thank A. Brydniak, G. Gillard, L. Laffon, C. Perrin, F. Sanchez and P. Vaast for their participation in data collection, and C. Rasse from the SMCS for advices concerning data analysis and interpretation. We thank the reviewers of our article for their very helpful comments. This article is number BRC 370 of the Biodiversity Research Center of the UCLouvain.

## Supplementary Material

**Table S1:**
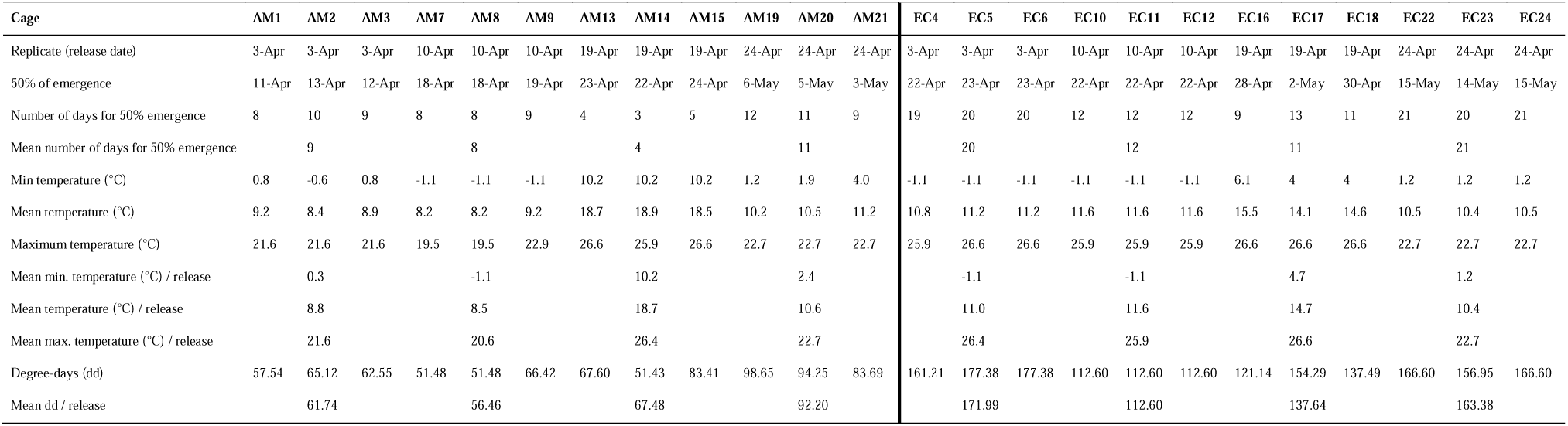
Date of emergence, mean temperatures and calculated degree-days for each cage containing *A. matricariae* (AM) *and E. cerasicola* (EC) mummies.

## Notes

### Competing Interest Statement

The authors have declared no competing interest.

### Summary of Updates

The author order was wrong

